## Supplementary Information and Figures for "Epistasis between mutator alleles contributes to germline mutation spectra variability in laboratory mice"

### Appendix 1

#### Missense mutations in *Setmar* are unlikely to contribute to epistasis in the BXDs

Unlike *Ogg1*, *Setmar* does not participate directly in base-excision repair, though its primate ortholog plays an indirect role in the repair of double-stranded DNA breaks via non-homologous end joining (NHEJ). In anthropoid primates, *SETMAR* encodes a fusion of two functional domains: a SET domain-containing histone methyltransferase and a transposase domain from the *Mariner* family (MAR) [84]; the mouse *Setmar* ortholog only encodes the histone methyltransferase domain. In human cell lines, *SETMAR* localizes to induced double-strand breaks (DSBs) and dimethylates nearby H3K36, which promotes the recruitment of DNA repair components involved in NHEJ to the DSB [37]. There is also evidence that overexpression of *SETMAR* (also known as *Metnase*) improves the efficiency of NHEJ [38] and leads to increased cell survival following exposure to ionizing radiation [38]. Point mutations in either the SET or MAR domains significantly reduced the ability of *SETMAR* to promote non-homologous end joining and DNA repair [38,39,40], suggesting that both domains are needed for its role in DNA repair. Another study found that overexpression of the isolated SET and MAR domains, but not of wild-type *SETMAR*, had a modest effect on NHEJ repair; overexpression of the SET domain slightly *decreased* NHEJ repair of a linearized plasmid in human cells, while overexpression of the *Mariner*-derived domain increased NHEJ relative to controls [85].

Taken together, these results suggest that both the SET and transposase domains of primate *SETMAR* are important for *SETMAR*-mediated DNA repair. The p.Leu103Phe missense mutation that differentiates C57BL/6J and DBA/2J (Table 1) resides within the *Setmar* pre-SET domain and occurs at an amino acid residue that is predicted to be deleterious by SIFT [47]. However, since the mouse *Setmar* ortholog lacks the *Mariner*-derived domain, we believe that the the p.Leu103Phe or p.Ser273Arg missense mutations are unlikely to affect C>A mutation rates in the BXDs. Moreover, we believe that the documented mutator phenotypes associated with *Ogg1*, as well as that gene's known role in base-excision repair, make it more likely candidate to underlie the epistatic interaction with *Mutyh* we observed in this study.

### Supplementary Figures

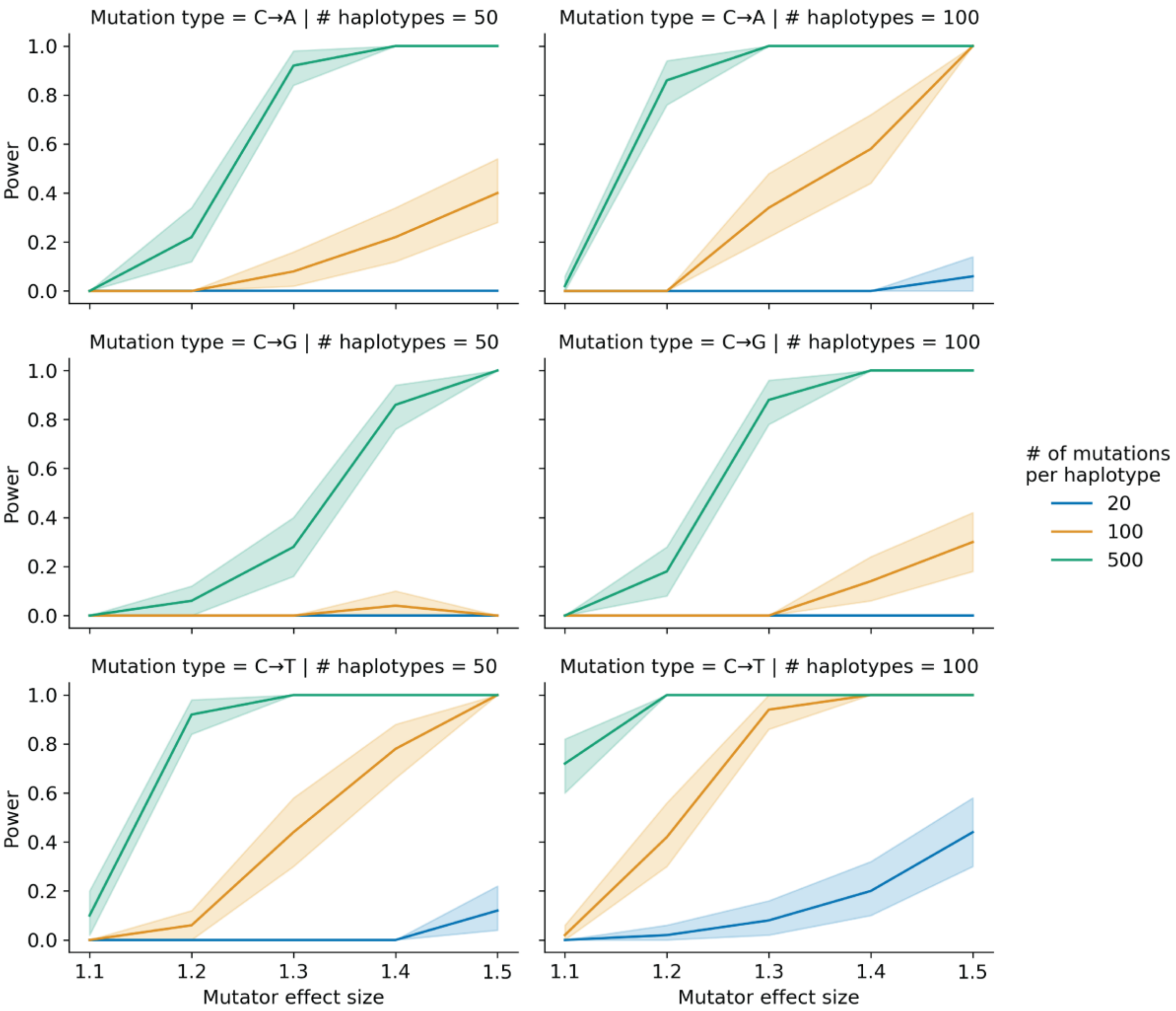


*Figure 1-figure supplement 1:* ***Simulations to assess the power of the aggregate mutation spectrum distance method.*** *In each of 50 trials, we simulated genotypes at 1,000 biallelic loci on a toy population of either 50 or 100 haplotypes as follows. At every locus on every haplotype, we drew a single floating point value from a uniform distribution* $[0,1)$*. If that value was less than or equal to 0.5, we set the allele to be "A"; otherwise, we set the allele to be "B". In each trial, we also simulated de novo germline mutations on the population of haplotypes, such that at a single locus* $g_i$*, we augmented the mutation rate of a particular* $k$*-mer by the specified effect size (an effect size of 1.5 indicates a 50% increase in the mutation rate) on haplotypes carrying "A" alleles. We then applied the aggregate mutation spectrum distance method to these simulated data and asked if the adjusted cosine distance at locus* $g_i$ *was greater than expected by chance. Given a specific combination of parameters, the y-axis denotes the fraction of 50 trials in which the simulated mutator allele could be detected at a significance threshold of p = 0.05. Shaded areas indicate the 95% bootstrap confidence interval surrounding that estimate.*

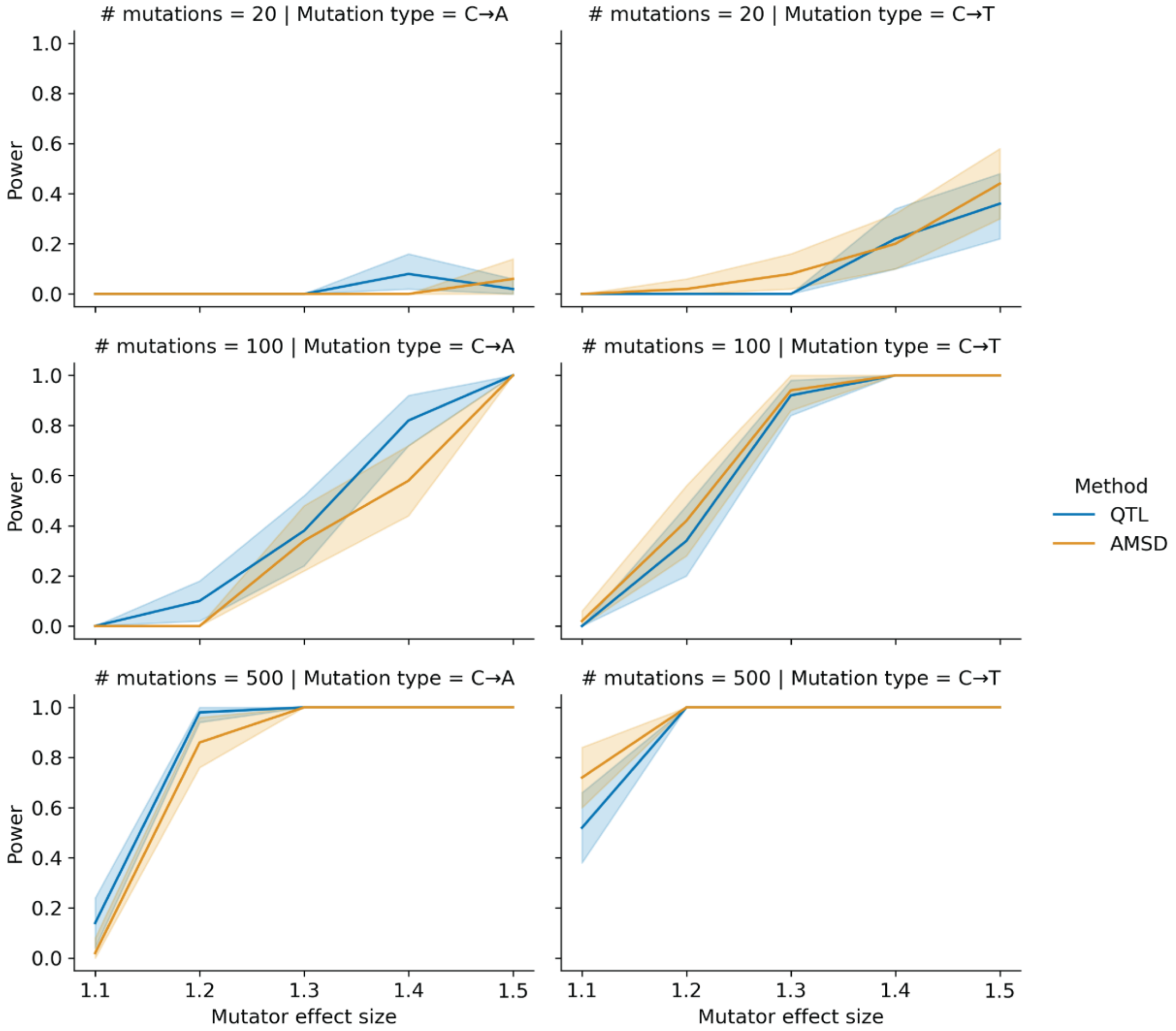


*Figure 1-figure supplement 2:* ***Comparing power between the aggregate mutation spectrum distance method and QTL mapping.*** *In each of 50 trials, we simulated genotypes at 1,000 biallelic loci on a toy population of 100 haplotypes as follows. At every locus on every haplotype, we drew a single floating point value from a uniform distribution* $[0,1)$. *If that value was less than or equal to 0.5, we set the allele to be "A"; otherwise, we set the allele to be "B". In each trial, we also simulated de novo germline mutations on the population of haplotypes, such that at a single locus* $g_i$, *we augmented the rate of the specified mutation type by the specified effect size (an effect size of 1.5 indicates a 50% increase in the mutation rate) on haplotypes carrying "A" alleles. We then applied the aggregate mutation spectrum distance method to these simulated data and asked if the adjusted cosine distance at locus* $g_i$ *was greater than expected by chance. Similarly, in each trial, we used R/qtl2 to perform a genome scan for QTL and asked if the log-odds score at* $g_i$ *was greater than expected by chance. Given a specific combination of parameters, the y-axis denotes the fraction of 50 trials in which the simulated mutator allele could be detected at a significance threshold of p = 0.05 (for AMSD) or at an alpha of* $\frac{0.05}{7}$ *(for*

*QTL mapping). Shaded areas indicate the 95% bootstrap confidence interval surrounding that estimate.*

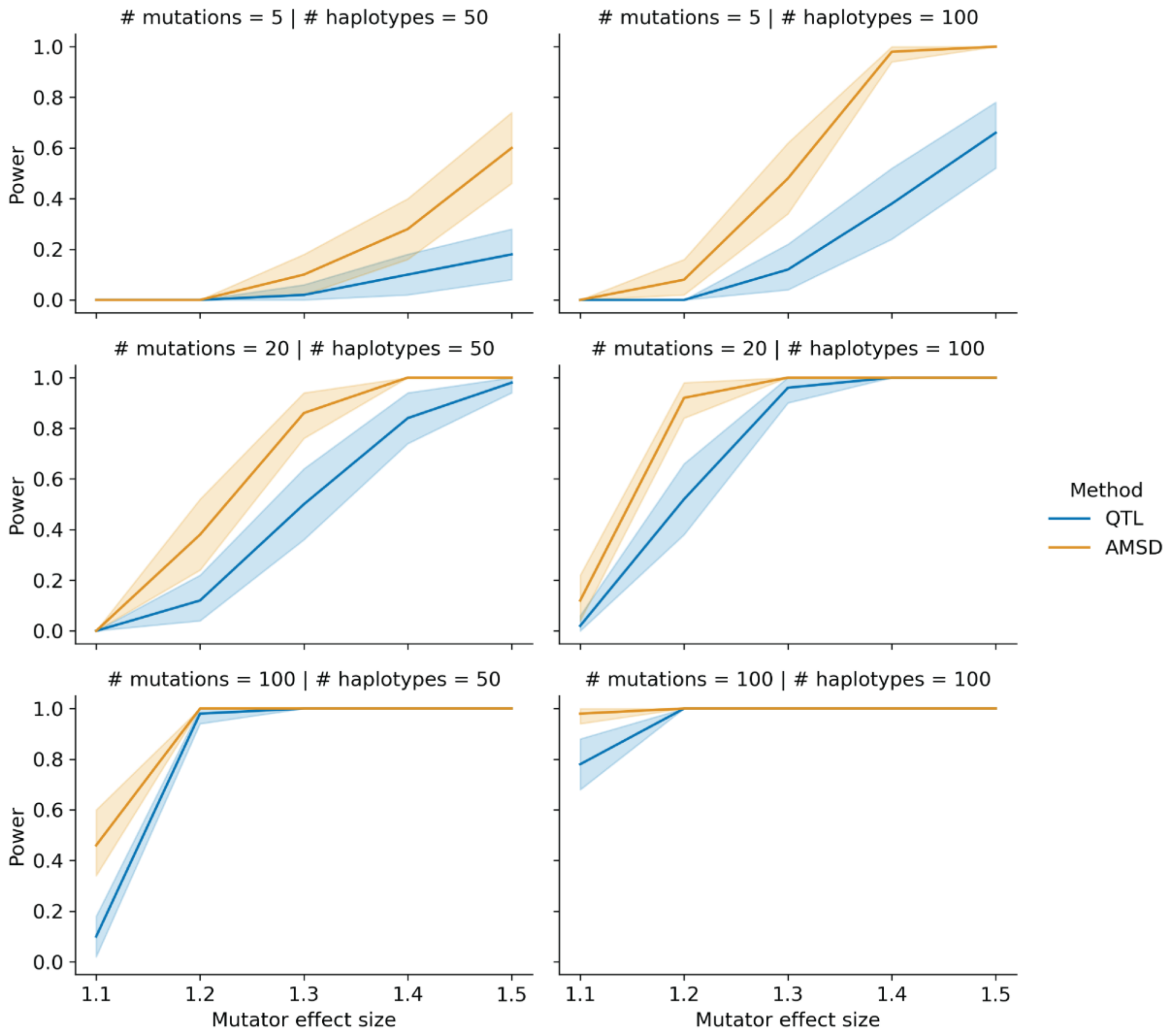


*Figure 1-figure supplement 3:* ***Comparing power between the aggregate mutation spectrum distance method and QTL mapping with variable counts of simulated mutations.*** *In each of 50 trials, we simulated genotypes at 1,000 biallelic loci on a toy population of 50 or 100 haplotypes as follows. At every locus on every haplotype, we drew a single floating point value from a uniform distribution* $[0,1)$*. If that value was less than or equal to 0.5, we set the allele to be "A"; otherwise, we set the allele to be "B". In each trial, we also simulated de novo germline mutations on the population of haplotypes, such that at a single locus* $g_i$*, we augmented the rate of the specified mutation type by the specified effect size (an effect size of 1.5 indicates a 50% increase in the mutation rate) on haplotypes carrying "A" alleles. To more closely approximate the BXD RILs, the mean number of simulated mutations on each haplotype was allowed to vary by a factor of 20 (see Materials and Methods for more details). We then applied the aggregate mutation spectrum distance method to these simulated data and asked if the adjusted cosine*

*distance at locus $g_i$ was greater than expected by chance. Similarly, in each trial, we used R/qtl2 to perform a genome scan for QTL and asked if the log-odds score at $g_i$ was greater than expected by chance. Given a specific combination of parameters, the y-axis denotes the fraction of 50 trials in which the simulated mutator allele could be detected at a significance threshold of p = 0.05 (for AMSD) or at an alpha of $\frac{0.05}{7}$ (for QTL mapping). Shaded areas indicate the 95% bootstrap confidence interval surrounding that estimate.*

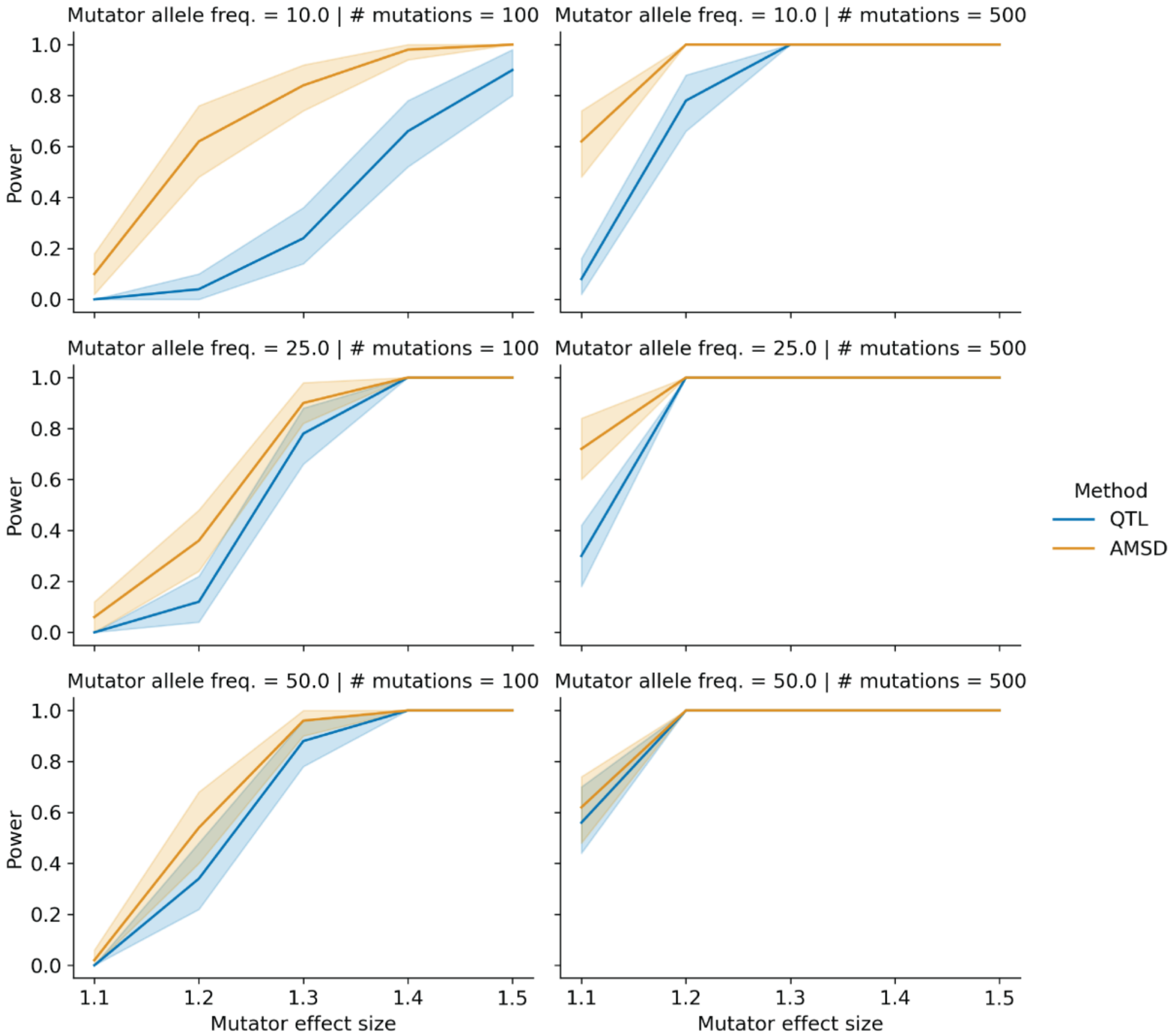


*Figure 1-figure supplement 4:* ***Comparing power between the aggregate mutation spectrum distance method and QTL mapping with variable mutator allele frequencies.*** *In each of 50 trials, we simulated genotypes at 1,000 biallelic loci on a toy population of 100 haplotypes as follows. At every locus on every haplotype (other than the simulated mutator locus, we drew a single floating point value from a uniform distribution $[0,1)$. If that value was less than or equal to 0.5, we set the allele to be "A"; otherwise, we set the allele to be "B". To model the effects of mutator allele frequencies on AMSD and QTL power, we allowed the expected frequency of "A"*

*alleles at the mutator allele marker to be either 0.1, 0.25, or 0.5 in these simulations. In each trial, we also simulated de novo germline mutations on the population of haplotypes, such that at a single locus* $g_i$*, we augmented the rate of the specified mutation type by the specified effect size (an effect size of 1.5 indicates a 50% increase in the mutation rate) on haplotypes carrying "A" alleles. We then applied the aggregate mutation spectrum distance method to these simulated data and asked if the adjusted cosine distance at locus* $g_i$ *was greater than expected by chance. Similarly, in each trial, we used R/qtl2 to perform a genome scan for QTL and asked if the log-odds score at* $g_i$ *was greater than expected by chance. Given a specific combination of parameters, the y-axis denotes the fraction of 50 trials in which the simulated mutator allele could be detected at a significance threshold of p = 0.05 (for AMSD) or at an alpha of* $\frac{0.05}{7}$ *(for QTL mapping). Shaded areas indicate the 95% bootstrap confidence interval surrounding that estimate.*

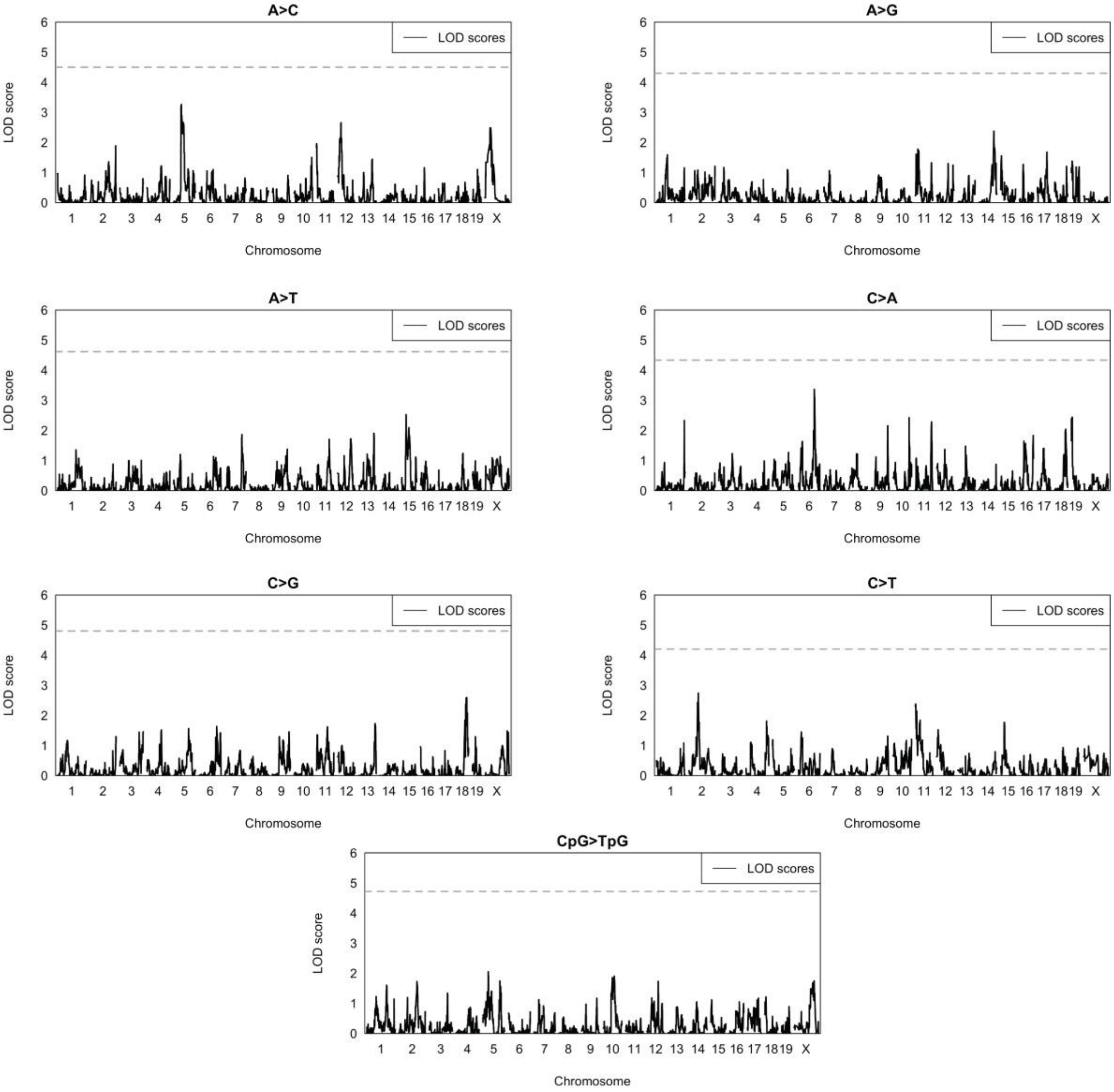


*Figure 2-figure supplement 1:* ***Quantitative trait locus scans for mutation spectrum phenotypes.*** *Using the BXDs with D genotypes at* `rs27509845` *(the marker with the highest cosine distance on chromosome 4; n = 66 BXDs, 42,171 total mutations), we used R/qtl2 to perform QTL scans for the fractions of each 1-mer mutation type. QTL scans also included a kinship matrix (that contained the pairwise genetic similarity between each pair of BXDs, calculated using the leave-one-chromosome-out method) as a random effect term using the* `kinship` *keyword argument in the* `scan1` *function. Plots show the log-odds (LOD) score at every genotyped marker in blue; the dotted black line represents the genome-wide LOD significance threshold (established using 1,000 permutations at an alpha of* $\frac{0.05}{7}$ *to account for the fact that 7 separate association tests were performed.)*

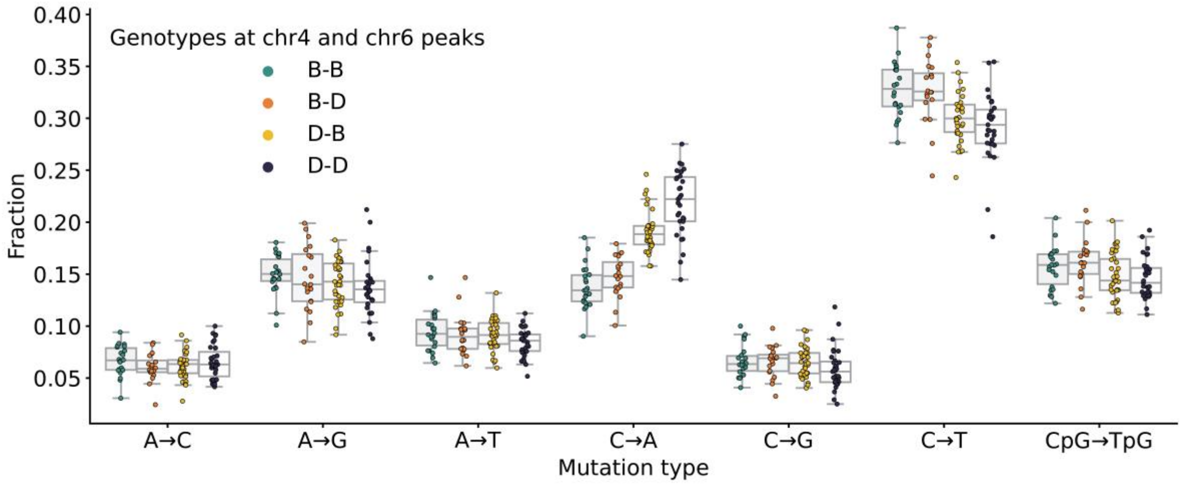


*Figure 3-figure supplement 1:* ***Mutation spectra comparison in BXD strains.*** *Fractions of de novo germline mutations in BXDs with either D or B genotypes at markers* `rs27509845` *and* `rs46276051`*, stratified by mutation type.*

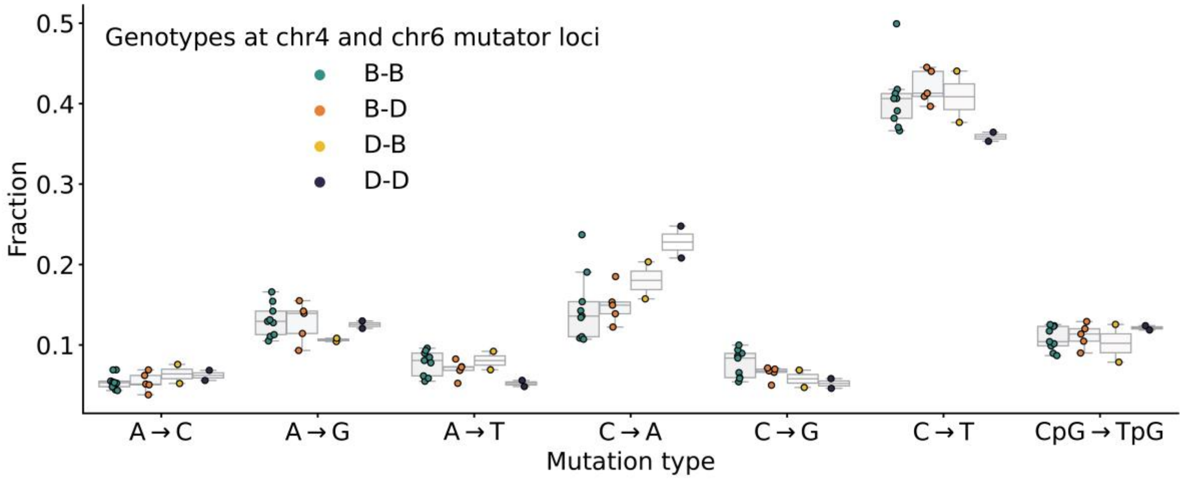


*Figure 3—figure supplement 2:* ***Mutation spectra comparison in Sanger Mouse Genomes Project strains.*** *Fractions of de novo germline mutations in Sanger MGP strains with either D or B haplotypes at the chromosome 4 and chromosome 6 mutator loci, stratified by mutation type.*

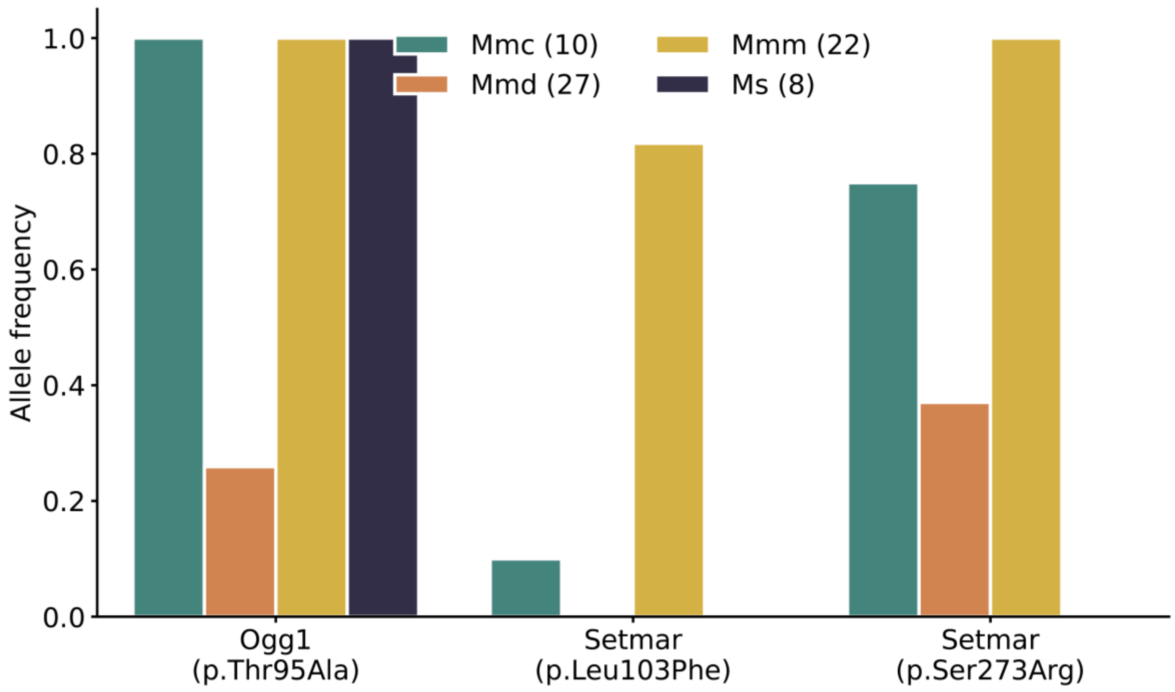


*Figure 3-figure supplement 3:* ***Frequency of nonsynonymous DNA repair mutations in wild mice.*** *Alternate allele frequencies of each nonsynonymous DNA repair mutation overlapping the chromosome 6 mutator locus were calculated in populations of wild-derived mice from Harr et al. [32]. Numbers of mice in each subpopulation are shown in parentheses. Mmc (Mus musculus castaneus), Mmd (Mus musculus domesticus), Mmm (Mus musculus musculus), and Ms (Mus spretus). The Mbd4 p.Asp129Asn mutation was not observed in any wild populations.*

### Supplementary Tables

*Table supplement 1: Significant cis-eQTLs for DNA repair genes in various tissues identified using GeneNetwork.*

| Gene name | Tissue name | # BXDs with expression data | Top significant marker | -log10(p) at top significant marker (GEMMA) | Additive effect of D allele on expression (GEMMA) |
|---|---|---|---|---|---|
| Ogg1 | Kidney | 53 | rsm10000004188 | 12.89 | -0.180 |
| Ogg1 | Liver | 50 | rsm10000004188 | 13.57 | -0.155 |
| Ogg1 | Spleen | 79 | rsm10000003418 | 4.73 | -0.056 |
| Ogg1 | Gastrointestinal | 46 | rs4173870 | 5.43 | -0.048 |
| Fancd2 | Gastrointestinal | 46 | rsm10000004199 | 8.60 | 0.133 |
| Ogg1 | Hippocampus | 67 | rsm10000004188 | 16.50 | -0.165 |
| Rad18 | Hippocampus | 67 | rsm10000003463 | 6.32 | 0.068 |

| Gene name | Tissue name | # BXDs with expression data | Top significant marker | -log10(p) at top significant marker (GEMMA) | Additive effect of D allele on expression (GEMMA) |
|---|---|---|---|---|---|
| Setmar | Hippocampus | 67 | rs13478947 | 11.03 | 0.141 |
| Mbd4 | Spleen | 79 | rsm10000004199 | 6.05 | 0.071 |

130
